# Negligible effects of read trimming on the accuracy of germline short variant calling in the human genome

**DOI:** 10.1101/2023.04.28.538608

**Authors:** Yury A. Barbitoff, Alexander V. Predeus

**Affiliations:** Institute of Bioinformatics Research and Education, Belgrade, Serbia

## Abstract

Next generation sequencing (NGS) has become a standard tool in the molecular diagnostics of Mendelian disease, and the precision of such diagnostics is greatly affected by the accuracy of variant calling from sequencing data. Recently, we have made a comprehensive evaluation of the performance of multiple variant calling pipelines, showing that state-of-the-art neural network-based methods show the best accuracy of variant discovery in the coding genome. In this work, we systematically evaluated the effects of adapters on the performance of variant calling tools using standard reference Genome-in-a-Bottle (GIAB) samples. We show that adapter trimming has no effect on the accuracy of the best-performing variant callers (e.g., DeepVariant) on whole-genome sequencing (WGS) data. For whole-exome sequencing (WES) datasets subtle improvement of accuracy was observed in some of the samples. In high-coverage WES data (∼200x mean coverage), adapter removal allowed for discovery of 2-4 additional true positive variants in only two out of seven datasets tested. Moreover, this effect was not dependent on the median insert size and proportion of adapter sequences in reads. Surprisingly, the effect of trimming on variant calling was reversed when moderate coverage (∼80-100x) WES data was used. Finally, we show that some of the recently developed machine learning-based variant callers demonstrate greater dependence on the presence of adapters in reads. Taken together, our results indicate that adapter removal is unnecessary when calling germline variants, but suggest that preprocessing methods should be carefully chosen when developing and using machine learning-based variant analysis methods.

## Introduction

Next-generation sequencing (NGS) has rapidly transformed the field of genetics and genomics (reviewed in Goodwin et al., 2016). Besides all of its applications in basic and applied research, NGS is quickly becoming a standard tool in clinical practice of medical geneticists (Biesecker et al., 2014). Despite the fact that NGS methods are widely used for rare disease diagnostics, the technology still has many important limitations. For example, commonly used second generation sequencing technologies based on short sequencing reads (e.g., Illumina or MGISEQ) are incapable of accessing certain regions of the human genome, such as regions with highly repetitive sequences and low mappability (Barbitoff et al., 2020; Ebbert et al., 2019). Our recent analysis showed that less than 90% of the coding genome sequence nucleotides could be considered high-confidence regions in which reliable variant calling is possible (Barbitoff et al., 2022). Furthermore, variant interpretation remains a major challenge, especially in the non-coding parts of the genome (Ellingford et al., 2022). Taken together, the aforementioned factors decrease the average rate of molecular diagnostics from NGS data (reviewed in Wright et al., 2018)..

Besides limitations associated with the technology itself, the choice of bioinformatic tools for data analysis may also greatly impact the diagnostic outcome. Recently, we have performed a systematic benchmark of state-of-the art variant calling pipelines (Barbitoff et al., 2022). The results suggested that most recent variant calling methods which make use of machine learning algorithms, such as DeepVariant (Poplin et al., 2018) perform substantially better compared to the widely used tools such as the Genome Analysis ToolKit (GATK) (McKenna et al., 2013; van der Auwera et al., 2014). Furthermore, we demonstrated that these solutions show no signs of being overfitted to the gold standard datasets. However, some machine learning (ML)-based approaches (e.g., GATK 2D CNN model or the Octopus random forest model) may produce puzzling results, for example, when facing substantial deviations in mean sequence coverage. These results suggest that careful choice of data preprocessing methods may become more critical for pipelines based on such variant callers.

Adapter trimming is widely considered as one of the routine steps of the variant calling pipeline; however, it is frequently omitted due to the ability of the most common read aligners to soft-clip adapter bases during mapping. A recent study in bacteria showed that read trimming may even be detrimental (Bush, 2020). The question of the impact of adapters on variant calling in the human genome is especially relevant given the aforementioned sensitivity of certain ML-based variant calling methods to various properties of the data. Hence, in this study we analyzed the effects of read trimming on variant calling in the human genome by evaluating the performance of different variant calling pipelines on trimmed and untrimmed data. We demonstrate that the effects of adapter removal on variant caller performance is negligible, and, despite certain performance gains for individual methods, does not affect the overall performance differences between pipelines observed in our previous work.

## Methods

### Data acquisition and preprocessing

We acquired the sequencing data and gold standard sets of variants for the Genome In A Bottle (GIAB) v. 4.2 release as described in our previous work (Barbitoff et al., 2022). For the downsampling experiment, we used the seqtk tool (https://github.com/lh3/seqtk) to sample 40% of read pairs for each individual WES sample. For adapter trimming, we used fastp v. 0.23.2 (Chen et al., 2018), Trimmomatic v. 0.39. (Bolger et al., 2014), and BBduk v. 37.62. Only built-in databases of Illumina adapters were used for trimming. Trimming was performed in the palindromic mode.

The proportions of adapter bases and adapter-containing reads were calculated using a set of custom shell scripts using the raw and trimmed FASTQ files. We considered that all bases removed during trimming correspond to adapters, and any read that has shortened due to trimming contained adapter sequences.

### Variant calling pipelines

Raw or preprocessed reads were aligned onto the GRCh37.p13 human reference genome assembly using the BWA MEM v. 0.7.17-r1188 read alignment software (Li and Durbin, 2011). Mapped reads were converted to BAM and sorted by coordinate using samtools; duplicate reads were marked using the GATK MarkDuplicates tool. The resulting BAM file was used for variant calling with six different software tools: Clair3 v. 1.0.0 (Luo et al., 2020), DeepVariant v. 1.4.0 (Poplin et al., 2018). Freebayes v. 1.3.1 (Garrison and Marth, 2012), Genome Analysis ToolKit HaplotypeCaller v. 4.3.0 (DePristo et al., 2011), Octopus v. 0.7.4 (Cooke et al., 2021), and Strelka2 v. 2.9.10 (Kim et al., 2018). Variants called by Octopus were additionally filtered using the built-in random forest model. For GATK HaplotypeCaller (GATK-HC), an additional step of base quality score recalibration (BQSR) was performed, and variants were called with GATK-HC in the single-sample mode using the recalibrated BAM file. Raw GATK-HC VCF files were filtered using either the hard filtering method (with the recommended settings) or the CNNScoreVariants (Friedman et al., 2020) model (only 1D model was used due to generally low robustness of the 2D model (Barbitoff et al., 2022)). For both CNN models, different tranche values were tested, and SNP tranche value of 99.9 and indel tranche value of 99.5 were used as showing the best performance. For Strelka2, an additional pipeline step was performed to generate a list of candidate indel sites using Manta (Chen et al., 2016). For all tools, variant calling was limited to a set of CDS intervals containing flanking (150 bp up- and downstream) regions.

### Benchmarking strategy

Benchmarking of variant calling results was performed using the hap.py toolkit (Krusche et al., 2019). Analysis was restricted to CDS intervals only. For WES data, an additional BED file was used to limit the evaluation to targeted regions of the exome in the Agilent SureSelect v7 exome capture kit. A common set of high-confidence variant calling intervals was used to make all analyses (the set of intervals was constructed by merging GIAB v. 4.2 high-confidence regions for all seven samples included in the study (Barbitoff et al., 2022)).

When evaluating the impact of read trimming on the variant calling accuracy, a value of each metric (precision, recall, F1 score) obtained using untrimmed data was subtracted from the corresponding metric value for the trimmed data. The number of affected variants was calculated by subtracting the number of true positive (TP) and false positive (FP) variants for the untrimmed data from the corresponding number obtained using the trimmed dataset.

### Data availability

All data and code pertinent to the analysis presented in this paper are available at: https://github.com/ibre-research/trimming-effects/

## Results

To assess the effects of adapter removal on the efficiency of variant discovery we collected gold-standard WES and WGS datasets for the seven samples from the GIAB consortium - NA12878 (CEU), the Chinese trio, and the Ashkenazi trio (Barbitoff et al., 2022; Supplementary Table 1). We began by evaluating the overall presence of adapter sequences in these datasets. To this end, we calculated the percentage of bases corresponding to adapter sequences, as well as the proportion of reads containing adapters (see Methods for details). The analysis showed that, expectedly, adapters are present in higher levels in WES compared to WGS samples (Figure 1a). We also observed a substantial variation in the fraction of adapter bases and reads containing adapters among the WES samples - for example, the proportion of reads containing adapters varied from 8.1% to 35.2% (Figure 1a).

**Figure 1.**
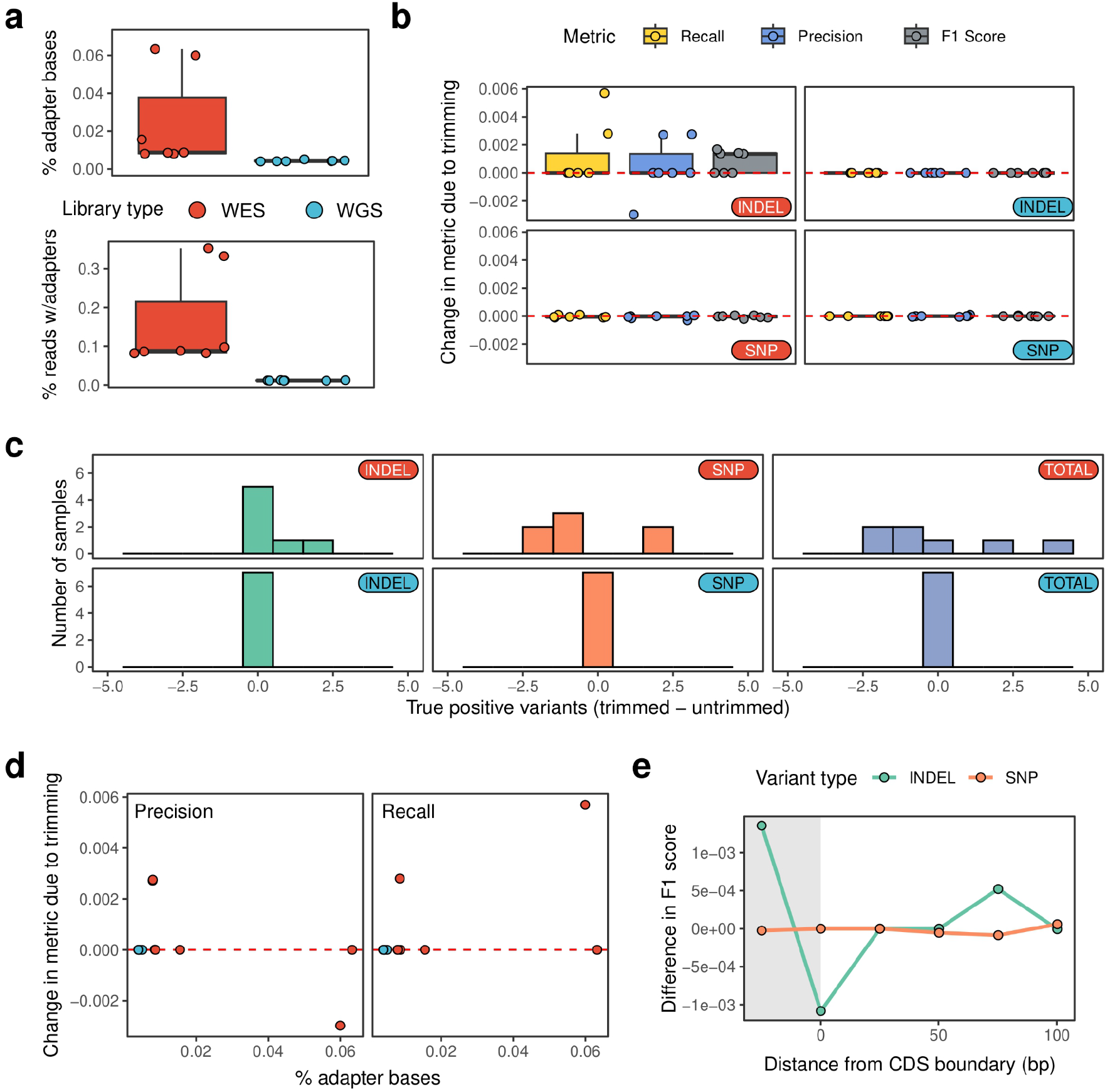
Adapter trimming has limited effects on variant calling accuracy. (a) The proportion of bases removed during adapter trimming with FASTP (Chen et al., 2018) (top) and the proportion of reads affected by this trimming (bottom) in WES and WGS data. (b) Boxplots of the differences in the benchmarking metrics (precision, recall, and F1 score) between trimmed and untrimmed data for SNPs and indels in WES and WGS. (c) A histogram showing the difference in the number of true positive variant calls (SNP, indel, and combined (denoted as “TOTAL”)) in WES (top) and WGS (bottom) data. (d) A scatterplot showing the correspondence between the fraction of adapter bases (shown in (a)) and the effect of adapter removal on precision (top) and recall (bottom). Only values for indel variants are shown. (e) Median difference in F1 score of SNP and indel discovery in WES data in 25bp-wide regions located at the indicated distance from the CDS boundary. Shaded area corresponds to the values observed for in-CDS variants.. In (b-e), shown are the results obtained using the variant calling pipeline which showed best performance in our recent benchmark (Barbitoff et al., 2022) (BWA MEM + DeepVariant).

We next went on to evaluate the impact of adapter trimming on the results of variant calling with the best-performing variant calling pipeline identified in our earlier benchmark - a combination of the BWA MEM aligner (Li and Durbin, 2011) and the DeepVariant (Poplin et al., 2018) variant caller (Barbitoff et al., 2022). To this end, we ran this pipeline for raw FASTQ data for all samples, as well as on the same data preprocessed using the FASTP method (Chen et al., 2018; 10.1093/bioinformatics/bty560). A comparison of benchmarking results (see Methods for details on benchmarking procedure) showed that adapter trimming had very limited effects on both precision and recall (Figure 1b). Consistently with the analysis of adapter proportion in reads, no performance difference was observed for the raw and trimmed WGS data, neither for SNP nor for indel variants. For WES data, no precision or recall gain was observed for the majority of the samples; however, a slight increase in precision and recall of indel calling was observed for the majority (four out of seven) datasets. Positive effects on recall, however, were observed in only two of the samples. Furthermore, when the total (SNPs and indels combined) number of true positive (TP) variants is considered, more TPs were discovered in two samples (two for HG002 and four - for HG006) (Figure 1c), while a decrease in the total TP count was observed for the majority (four out of seven) datasets.

We further questioned whether the observed subtle performance gains for the trimmed data correlate with the proportion of adapters in reads. To answer this question, we plotted the difference in both precision and recall (Figure 1d) observed for indels in WES samples against the proportion of adapter bases. To our surprise, we observed no direct dependence between an increase and precision or recall and the total adapter content (Figure 1c-d). Given this result, we hypothesized that the difference in variant calling accuracy may be driven by variants located in adapter-enriched regions located at the exon boundaries in WES. To test this hypothesis, we evaluated the performance of DeepVariant in regions located in the vicinity of the exon boundary. Contrary to our expectations, we observed a reverse effect of adapter trimming on variant calling outcome in the immediate exon vicinity (0-25 bp) (Figure 1e). In regions located further away from the exon margin, trimming had no median effect on the F1 scores.

Given these results, we next asked if a more reproducible effect of trimming on variant calling accuracy can be observed when using other bioinformatic tools for adapter removal. To this end, we applied two other commonly used methods for trimming adapters - Trimmomatic (Bolger et al., 2014) and BBDUK (https://github.com/BioInfoTools/BBMap). Evaluation of variant calling results showed that all tools produce similar results, with only a slightly lower precision when using Trimmomatic (Supplementary Figure S1).

At the next stage of our analysis, we decided to compare the effects of adapter trimming on other variant calling methods. For this comparison, we used a set of nine variant calling and filtering methods from our previous analysis (Barbitoff et al., 2022). We applied this set of tools to the set of GIAB datasets, focusing solely on WES samples due to markedly higher adapter content. Analysis of variant calling results showed that some of the tools demonstrate greater sensitivity to presence of adapters in reads. In particular, Clair3 and Octopus (with the random forest filtering model) showed a significant and reproducible performance increase on trimmed data compared to untrimmed (Figure 2a). Furthermore, Strelka2 showed a large increase in performance for samples with high adapter content, though the median difference in F1 score across all samples was subtle. Importantly, the observed gains in the F1 score were not sufficient for any tool to surpass DeepVariant when comparing the F1 scores on trimmed data (Figure 2b).

**Figure 2.**
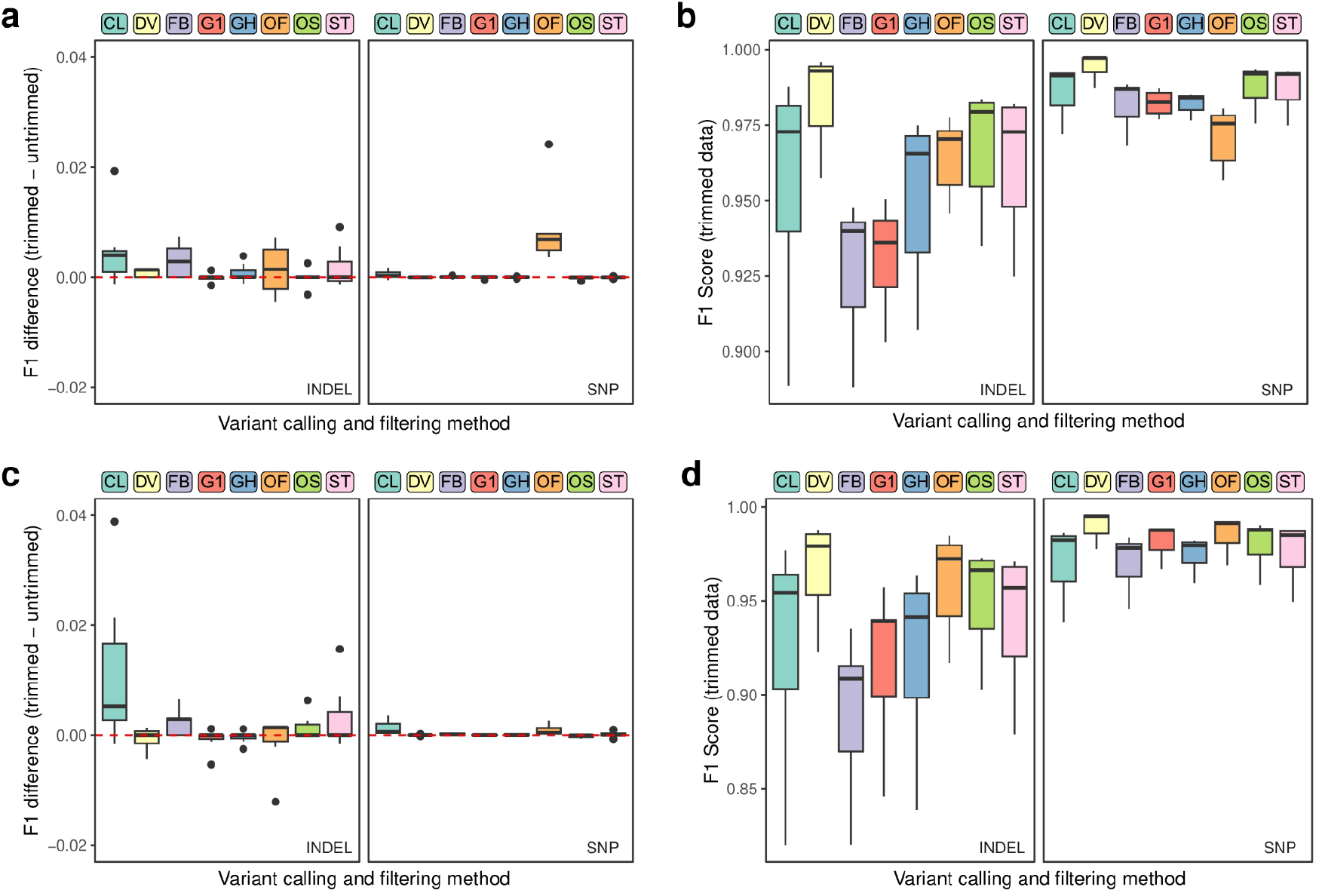
Some of the variant calling methods demonstrate greater sensitivity to the presence of adapters in reads. (a) Boxplots showing the differences in F1 score values between variant calls produced on trimmed and untrimmed high coverage WES data using the indicated variant calling pipelines. (b) F1 scores of indel and SNP calling with the indicated pipelines on FASTP-trimmed of high-coverage WES data. (c, d) Same as in (a, b), but for moderate(∼70-100x) coverage datasets. CL - Clair3, DV - DeepVariant, FB - Freebayes, G1 and GH - GATK HaplotypeCaller with 1D CNN or hard filtering, OF and OS - Octopus with random forest or standard filtering, ST - Strelka2.

Finally, we hypothesized that the low magnitude of effect of adapter trimming on variant calling accuracy may be explained by the high coverage of the WES datasets used for analysis (the mean coverage of CDS bases was more than 200x). To evaluate this hypothesis, we downsampled 40% of the reads in each sample to obtain a set of WES data with substantially lower mean coverage (∼80x). Such WES datasets better represent a typical exome sample. After generating the downsampled WES data, we applied all tested variant calling pipelines to this set of samples, and evaluated the accuracy of the resulting variant calls. The analysis showed that the effects of adapter trimming for downsampled data were on average less pronounced compared to the ones observed using high-coverage samples (Figure 2c). The effect of trimming for the best-performing pipeline based on DeepVariant was less pronounced in downsampled compared to original data. Similarly, Octopus with the random forest filtering mode also showed a substantially lower impact of trimming on moderate-coverage datasets compared to high-coverage ones. For GATK-HC and Strelka2 the results were similar on high- and moderate-coverage data, while Octopus with a standard filtering scheme showed an irreproducible positive impact of trimming on indel calling (Figure 2a). The only exception from the general trend was Clair3 which showed markedly greater performance gains upon trimming in moderate-coverage WES datasets. We also compared the performance of each individual pipeline on high-coverage and low-coverage WES data. In concordance with the results of our previous analysis (Barbitoff et al., 2022), only Octopus with random forest filtering model performed better on low-coverage compared to high-coverage datasets (Supplementary Figure S2); nevertheless, DeepVariant remained the best-performing solution for the downsampled data both before and after adapter trimming (Figure 2d)

## Discussion

Pre-processing and trimming of raw sequencing reads is commonly considered a standard step in virtually any data analysis workflow. However, the development of both laboratory protocols for sequencing library preparation and bioinformatic tools for the analysis reopens the question of the necessity of read trimming procedure for accurate analysis and interpretation of sequencing results. A recent analysis in bacteria (Bush, 2020) suggested that read trimming has minimal effects on the accuracy of variant calling. Given this report, as well as the result of our recent benchmark of state-of-the-art variant callers, we conducted a systematic analysis of the impact of read trimming on the accuracy of germline variant analysis in the human genome.

Concordantly with the results of (Bush, 2020), we observed a very limited effect of adapter removal on the accuracy of variant calling. For the majority of both WGS and WES samples, the difference in the performance of the most accurate combination of tools (the BWA MEM aligner and the DeepVariant variant caller) between trimmed and untrimmed data was equal to zero for both SNP and indel variants. Positive effects of adapter removal on variant calling were observed only for several variant sites in some of the WES samples. This result is concordant with the overall higher fraction of adapter bases in WES, and corroborates our previous findings regarding the major role of insert size in defining the limits of CDS coverage for WES libraries (Barbitoff et al., 2020).

According to our previous analysis, variant calling in the CDS-flanking regions is the main driver of lower variant calling accuracy for WES data (Barbitoff et al., 2022). This observation suggests that a positive effect of adapter removal on variant discovery from WES data may be driven by variants in such exon boundary regions, which should be enriched for adapters. However, our assessment of the impact of adapter removal on variant calling in the immediate vicinity of the CDS does not support this hypothesis (Figure 1e).

Among the combinations of variant calling and filtering methods tested, Clair3 and Octopus showed a significant dependence on the presence of adapters in reads (Figure 2a, c). For Octopus, this dependence was reproducibly observed only for the random forest-based filtering. Furthermore, our results confirm that a decrease in coverage leads to an unexpected improvement in the accuracy of variant filtering with this model (Supplementary Figure 2). These results are consistent with the greater dependence of the random forest filtering method in Octopus on coverage and other sequence properties (Barbitoff et al., 2022). Similarly, greater impact of adapter removal on Clair3 performance is consistent with its substantial dependence on alignment properties (Barbitoff et al., 2022). It is also important to note that our comparison of the accuracy of different pipelines on trimmed data confirms that DeepVariant remains the best-performing solution for germline variant analysis (Figure 2b, d). Taken together, the aforementioned findings provide additional support to the results of our earlier benchmark, and emphasize the necessity of thorough testing and validation of ML-based variant calling and filtering solutions.

Besides the overall negligible impact of adapter trimming on variant calling accuracy, adapter removal has other important consequences which affect the necessity of this procedure. First, removing adapter sequences from the reads leads to data loss if the raw read files are not stored separately. While the adapter sequences by themself do not contain any relevant information, their absence may affect future re-analysis of the data, including independent quality assessment. On the other hand, presence of adapter sequences in reads may complicate the visual inspection of the alignment results, especially for inexperienced users. In addition, both presence or absence of adapters may affect certain types of post-analysis of the alignment files.

It is also important to emphasize that our observations are valid for germline variant calling. In case of somatic variant discovery, which is known to have a much greater sensitivity to errors in reads and requires a different approach to data processing (Koboldt, 2020). Given the aforementioned features of somatic variant calling, it is possible to suggest that adapter trimming will have a much greater positive impact in this case.

Overall, we believe that our findings argue against adapter removal for germline variant calling in the human genome, While positive effects of read trimming can be observed only for some of the WES samples, trimming does not provide any noticeable performance gain in neither WGS nor the majority of WES datasets, and may even decrease the accuracy of analysis..

## Acknowledgements

We thank JetBrains Ltd. for providing financial support and computing resources for the project. The authors declare no conflict of interest.

## References

1. Barbitoff, Y. A., Abasov, R., Tvorogova, V. E., Glotov, A. S. & Predeus, A. V. Systematic benchmark of state-of-the-art variant calling pipelines identifies major factors affecting accuracy of coding sequence variant discovery. BMC Genomics 23, 1–17 (2022).

2. Barbitoff, Y. A. et al. Systematic dissection of biases in whole-exome and wholegenome sequencing reveals major determinants of coding sequence coverage. Sci. Rep. 10, 1–13 (2020).

3. Biesecker, L. G. & Green, R. C. Diagnostic Clinical Genome and Exome Sequencing. N. Engl. J. Med. 370, 2418–2425 (2014).

4. Bolger, A. M., Lohse, M. & Usadel, B. Trimmomatic: A flexible trimmer for Illumina sequence data. Bioinformatics 30, 2114–2120 (2014).

5. Bush, S. J. Read trimming has minimal effect on bacterial SNP-calling accuracy. Microb. Genomics 6, 1–13 (2020).

6. Chen, S., Zhou, Y., Chen, Y. & Gu, J. Fastp: An ultra-fast all-in-one FASTQ preprocessor. Bioinformatics 34, i884–i890 (2018).

7. Cooke, D. P., Wedge, D. C. & Lunter, G. A unified haplotype-based method for accurate and comprehensive variant calling. Nat. Biotechnol. (2021) doi:10.1038/s41587-021-00861-3.

8. DePristo, M. A. et al. A framework for variation discovery and genotyping using nextgeneration DNA sequencing data. Nat. Genet. 43, 491–498 (2011).

9. Ebbert, M. T. W. et al. Systematic analysis of dark and camouflaged genes reveals disease-relevant genes hiding in plain sight. Genome Biol. 20, 97 (2019).

10. Ellingford, J. M. et al. Recommendations for clinical interpretation of variants found in non-coding regions of the genome. Genome Med. 14, 1–19 (2022).

11. Friedman, S., Gauthier, L., Farjoun, Y. & Banks, E. Lean and deep models for more accurate filtering of SNP and INDEL variant calls. Bioinformatics 36, 2060–2067 (2020).

12. Garrison, E. & Marth, G. Haplotype-based variant detection from short-read sequencing. aRxiv 1–9 (2012).

13. Goodwin, S., McPherson, J. D. & McCombie, W. R. Coming of age: Ten years of next-generation sequencing technologies. Nat. Rev. Genet. 17, 333–351 (2016).

14. Koboldt, D. C. Best practices for variant calling in clinical sequencing. Genome Med. 12, 1–13 (2020).

15. Krusche, P. et al. Best practices for benchmarking germline small-variant calls in human genomes. Nat. Biotechnol. 37, 555–560 (2019).

16. Li, H. & Durbin, R. Fast and accurate short read alignment with Burrows-Wheeler transform. Bioinformatics 25, 1754–1760 (2009).

17. Luo, R. et al. Exploring the limit of using a deep neural network on pileup data for germline variant calling. Nat. Mach. Intell. 2, 220–227 (2020).

18. McKenna, A. et al. The Genome Analysis Toolkit: A MapReduce framework for analyzing next-generation DNA sequencing data. Genome Res. 10, 1297–1303 (2010).

19. Poplin, R. et al. A universal snp and small-indel variant caller using deep neural networks. Nat. Biotechnol. 36, 983 (2018).

20. Van der Auwera, G. A. et al. From FastQ Data to High-Confidence Variant Calls: The Genome Analysis Toolkit Best Practices Pipeline. Curr. Protoc. Bioinforma. 10.1-10.33 (2013) doi:10.1002/0471250953.bi1110s43.

21. Wright, C. F., FitzPatrick, D. R. & Firth, H. V. Paediatric genomics: diagnosing rare disease in children. Nat. Rev. Genet. 19, 253–268 (2018).

22. Zook, J. M. et al. Extensive sequencing of seven human genomes to characterize benchmark reference materials. Sci. data 3, 160025 (2016).

