## Supplementary Figures for "Negligible effects of read trimming on the accuracy of germline short variant calling in the human genome"

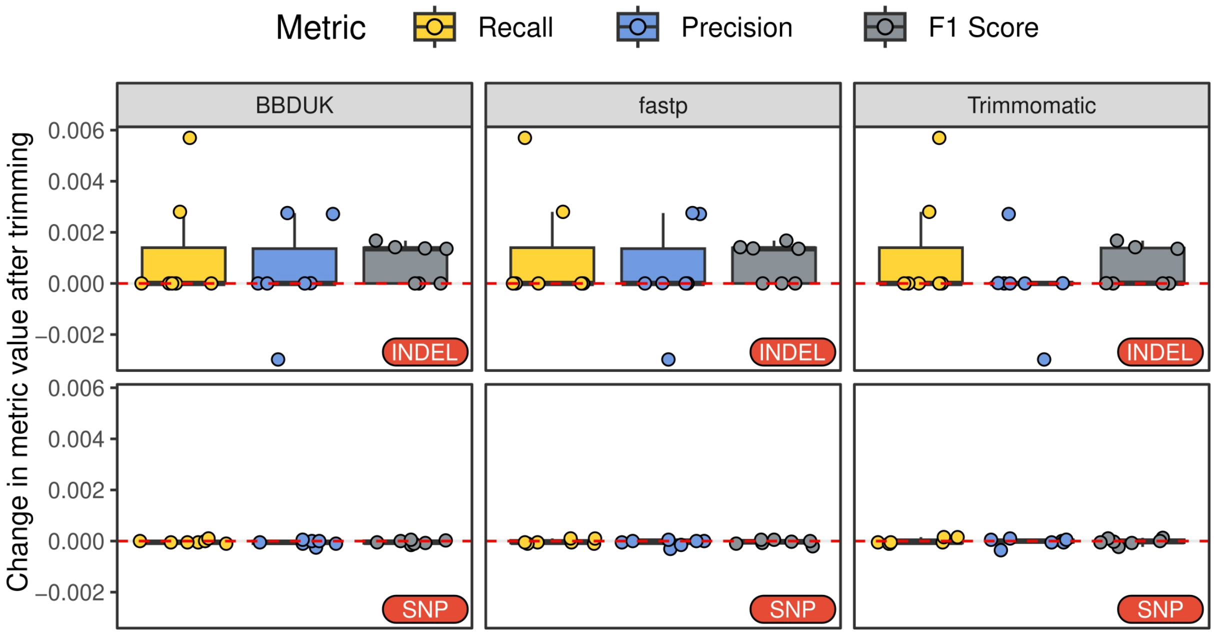


**Figure S1.** The effects of adapter removal with the indicated read trimming software. Shown are differences in metric values (precision, recall, and F1 score) between trimmed and untrimmed GIAB WES data.

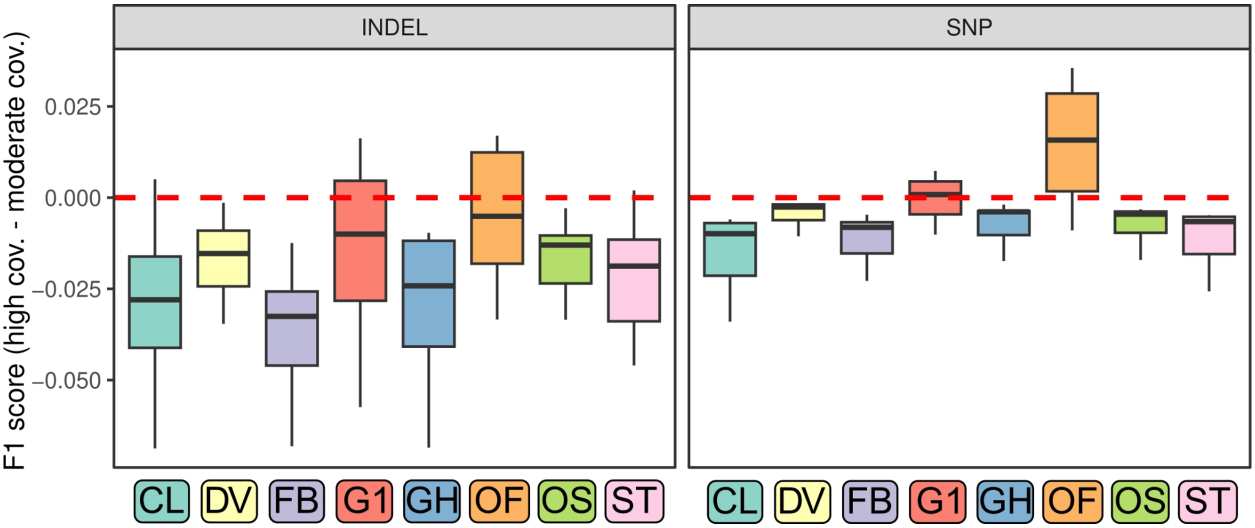


**Figure S2.** The difference in the F1 scores of SNP and indel calling with indicated variant calling pipelines on FASTP-trimmed high- and moderate-coverage WES datasets. CL - Clair3, DV - DeepVariant, FB - Freebayes, G1 and GH - GATK HaplotypeCaller with 1D CNN or hard filtering, OF and OS - Octopus with random forest or standard filtering, ST - Strelka2.
